# Incorporation of the HIV-1 envelope glycoprotein into viral particles is regulated by the tubular recycling endosome in a cell type-specific manner

**DOI:** 10.1101/2023.12.17.572063

**Authors:** Grigoriy Lerner, Lingmei Ding, Kathleen Candor, Paul Spearman

## Abstract

The HIV-1 envelope glycoprotein (Env) is incorporated into particles during assembly on the plasma membrane (PM). Env initially reaches the PM through the secretory pathway, after which it is rapidly endocytosed via an AP-2- and clathrin-dependent mechanism. Here we show that endocytosed cell surface Env enters the tubular recycling endosome compartment (TRE). Trafficking to the TRE was dependent upon motifs within the CT previously implicated in Env recycling and particle incorporation. Depletion of TRE components MICAL-L1 or EHD1 led to defects in Env incorporation, particle infectivity, and viral replication. Remarkably, defects were limited to cell types defined as nonpermissive for incorporation of CT-deleted Env, including monocyte-derived macrophages, and not observed in 293T, HeLa, or MT-4 cells. This work identifies the TRE as an essential component of Env trafficking and particle incorporation, and provides evidence that the cell type-dependent incorporation of Env is defined by interactions with components of the TRE.

## INTRODUCTION

Incorporation of the HIV-1 envelope glycoprotein (Env) into budding HIV virions is essential for the production of infectious viral progeny. The HIV-1 Gag protein is synthesized on cytosolic ribosomes, and travels to the particle assembly site on the PM through a poorly-defined mechanism. Env is synthesized as the gp160 precursor protein on ER-bound ribosomes, where it trimerizes and undergoes initial steps of glycosylation.^1,2^ Env trimers then traffic to the Golgi apparatus, where additional glycan modifications and cleavage by furin-like proteases take place, leading to the mature gp120/gp41 heterotrimer that is transported to the PM.^3,4^ Upon arrival at the PM, Env trimers are rapidly endocytosed and delivered to internal endosomal compartments in a clathrin- and AP2-dependent manner.^5,6^ How endocytosed Env returns to the PM for incorporation into budding virions during the assembly process remains incompletely understood.

The Env cytoplasmic tail (CT) plays a key role in directing cellular trafficking and particle incorporation of Env. The CT contains a myriad of trafficking motifs, including the membrane proximal YXXΦ clathrin binding motif,^6–8^ multiple dileucine motifs involved in binding to clathrin adaptor proteins,^9,10^ and retromer-interacting sequences (also called inhibitory sequences, IS1 and IS2).^11,12^ The N-terminal portion of the lentiviral lytic peptide 3 (LLP3) region of the CT contains tyrosine- and tryptophan-based motifs that regulate Env incorporation as indicated by loss of incorporation upon their deletion or mutagenesis.^13–16^. Remarkably, however, an intact CT is not required for particle incorporation when particles are produced in some cell types, including 293T and MT4 cells, which are termed “permissive” for incorporation of CT-deleted Env.^17^ Other cell types, including H9, CEM, and Jurkat T cell lines incorporated Env efficiently only in the presence of an intact CT, and are termed “nonpermissive” for incorporation of CT-deleted Env. Env incorporation and replication of HIV-1 in primary T cells and macrophages requires an intact CT, indicating that the nonpermissive phenotype is dominant in those cell types that are most relevant for HIV-1 replication. To help explain the crucial role played by the CT in Env trafficking, we have proposed that cell type specificity may be defined by differences in host trafficking pathways between permissive and nonpermissive cell types.^18,19^

We previously described a role for the Rab-related adaptor protein Rab11-FIP1C in regulating Env incorporation into particles.^14,19,20^ Interventions that disrupted FIP1C or that led to condensation and compromised function of the endosomal recycling compartment (ERC) significantly reduced Env incorporation in relevant cell lines such as the nonpermissive H9 cell line. These results identified FIP1C is a candidate recycling factor that regulates Env incorporation in a cell type-dependent manner. However, results with depletion of FIP1C were only partly consistent with this model, and a knockout strategy in primary CD4+ T cells did not prevent Env incorporation or viral replication,^21^ indicating that differences in expression or function of FIP1C alone do not explain cell type-dependence.^21^ Despite this, our findings with FIP1C provided an important connection between host recycling factors and Env incorporation that remains valid in many cell types, and support a model in which host recycling pathways determine CT-dependent Env incorporation into particles. Herein we identify the tubular recycling endosome (TRE) and its constituent factors as essential contributors to CT-dependent incorporation of Env.

The TRE consists of a network of tubular membranes extending from the ERC toward the plasma membrane that is implicated in the active segregation and recycling of host glycoproteins.^22–24^ The TRE is regulated by specific Rab proteins including Rab10 and Rab8, and characterized by the scaffold protein MICAL-like protein 1 (MICAL-L1) and F-BAR protein Syndapin2 that recruit additional factors involved in glycoprotein sorting and recycling, including the ATPase EH domain containing 1 (EHD1) involved in membrane scission.^23,25^ Here we identify for the first time the trafficking of HIV-1 Env to the TRE, where it strongly colocalized with TRE constituents including MICAL-L1, EHD1, and Rab10. Localization to the TRE required an intact CT, and Env CT mutants previously shown to be defective in particle incorporation failed to reach the TRE. Most remarkably, we observed that depletion of TRE components MICAL-L1 or EHD1 resulted in Env incorporation defects in nonpermissive cell types including primary macrophages, indicating a link between cell type-specific incorporation of Env and TRE-dependent recycling. These findings are the first to elucidate a role for the TRE in HIV replication, and provide strong support for the importance of CT-dependent recycling in Env incorporation in relevant cells including primary cells.

## RESULTS

### HIV-1 Env is rapidly delivered to Tubular Recycling Endosomes enriched in PI(4,5)P2

We previously described a method of tagging the Env ectodomain using a fluorogen-activated peptide (FAP) tag inserted into the V2/V3 loop of gp120, allowing pulse-labeling of Env on the surface of the cell with a cell impermeant dye.^26^ Utilizing pulse-labeling of Env and live cell TIRF microscopy, we observed that Env was rapidly delivered into tubules underlying the plasma membrane (Figure 1A). The tubules were enriched in phosphatidylinositol (4, 5) bisphosphate (PIP2) as indicated by a GFP-tagged biomarker for PIP2, the pleckstrin homology domain of phospholipase C-δ1 (PLCδ1-PH), (Figure 1A and supplemental movie S1). We were surprised to see Env enriched on long tubular membranes, as Env stained in fixed cells following usual fixation practices does not show this pattern. However, we noted that preservation of tubular membranes requires fixation methods that avoid disruption of these somewhat fragile structures, which can be achieved by using pre-warmed formaldehyde fixation combined with very low concentrations of non-ionic detergents for permeabilization.^27,28^ Utilizing this modification of specimen preparation, we readily observed HIV-1 Env in tubular structures deeper in the cell, extending from a perinuclear location toward the periphery of the cell (Figure 1B). Reasoning that PIP2 has been shown to be enriched on lipid tubules of the TRE,^23^ we next asked if tubular Env localized with three additional characteristic markers of the TRE: MICAL-L1, EHD1, and Rab10. Indeed, Env was found in tubular structures showing significant overlap with markers of the TRE, including MICAL-L1, EHD1, and Rab10 (Figure 1B). To better define the Env-enriched tubules, we utilized structured illumination microscopy and measured diameters of tubules in the X-Y plane (Figure 1C). Measurements were taken from multiple cross-sections of multiple images such as the one shown in Figure 1C, and indicated a mean width of 177± 45 nm, consistent with the established diameter of the TRE.^29^ These findings confirm the presence of Env on an extended tubular network consistent with the TRE.

**Figure 1.**
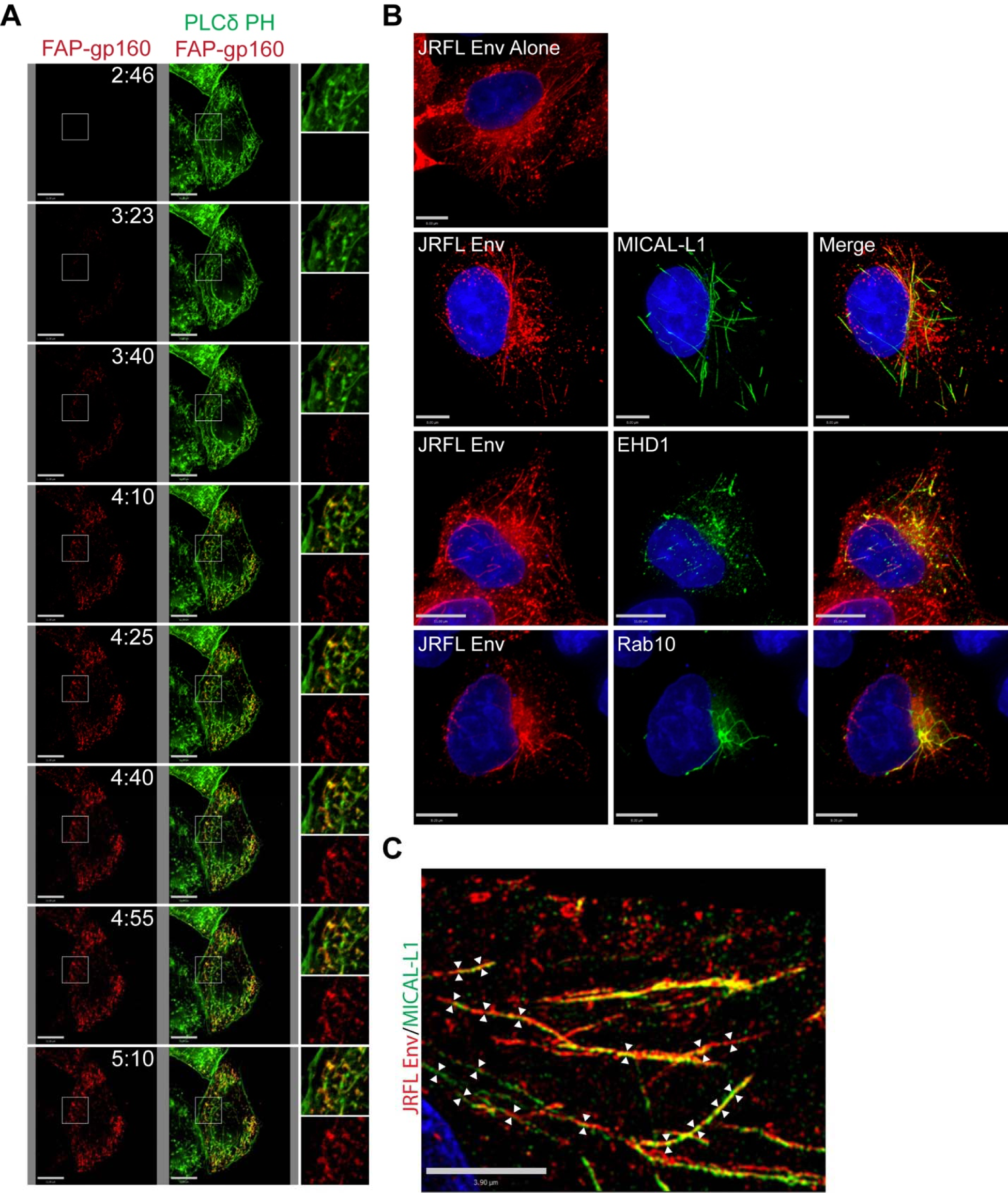
Endocytosed Env is present in elongated lipid tubules consistent with the TRE. (A) FAP-tagged Env was coexpressed in HeLa cells with GFP-tagged PLCδ-PH. Pulse-chase analysis was then performed using a non-membrane permeable fluorogen at 37°C, with Images captured at 15 second intervals. Images from the first 5 minutes of acquisition are shown here; see also Supplemental Video 1 for the full 20-minute imaging period. (B) Env expressed in HeLa cells was stained and imaged following fixation with pre-warmed paraformaldehyde and permeabilization with low amounts of nonionic detergent. Env alone was observed in tubular form under these conditions (top). Env was then examined in cells co-stained for characteristic TRE markers MICAL-L1, EHD1, and Rab10 (bottom panels). (C) Tubular structures showing dual staining for Env and MICAL-L1 were analyzed using structured illumination microscopy, and diameters measured at multiple points along the tubules (arrows).

### The Env CT is required for trafficking to the TRE

We next sought to examine the role of the Env CT in TRE localization. To do this, we employed several measures to evaluate colocalization of Env with an intact CT and compared them to Env lacking 144 C-terminal residues of the CT (CT144 Env). Full-length Env colocalized with TRE markers MICAL-L1, EHD1, and Rab10 as previously observed (Figure 2A). A region of interest was chosen in each of these images and is shown for each channel on the right (Figure 2A). Linear intensity profiles were drawn to bisect multiple tubules within the region of interest in order to further discern colocalized signal between Env and the markers of the TRE. In WT Env expressing cells, peaks of TRE marker fluorescence for MICAL-L1, EHD1, and Rab10 (green intensity plots, Figure 2B) correlated strongly with increased Env signal (red intensity plots, Figure 2B). In contrast, CT144 Env was not observed on tubular membranes, and failed to localize with tubules marked by MICAL-L1, EHD1, or Rab10 (Figure 2C). Linear intensity plots drawn perpendicular to tubules with TRE markers did not show any corresponding peaks of CT144 Env intensity (Figure 2D). Colocalization was next quantified for WT and CT144 Env with each of the three TRE markers using Manders’ colocalization coefficient (representing pixels positive for both Env and the respective tubule marker as a fraction of the total pixels of the tubule marker). This measurement verified the significant difference between WT and CT144 Env in colocalization with TRE markers (Figure 2E). Taken together, these data indicate that the enrichment of Env on TRE membranes requires an intact CT.

**Figure 2.**
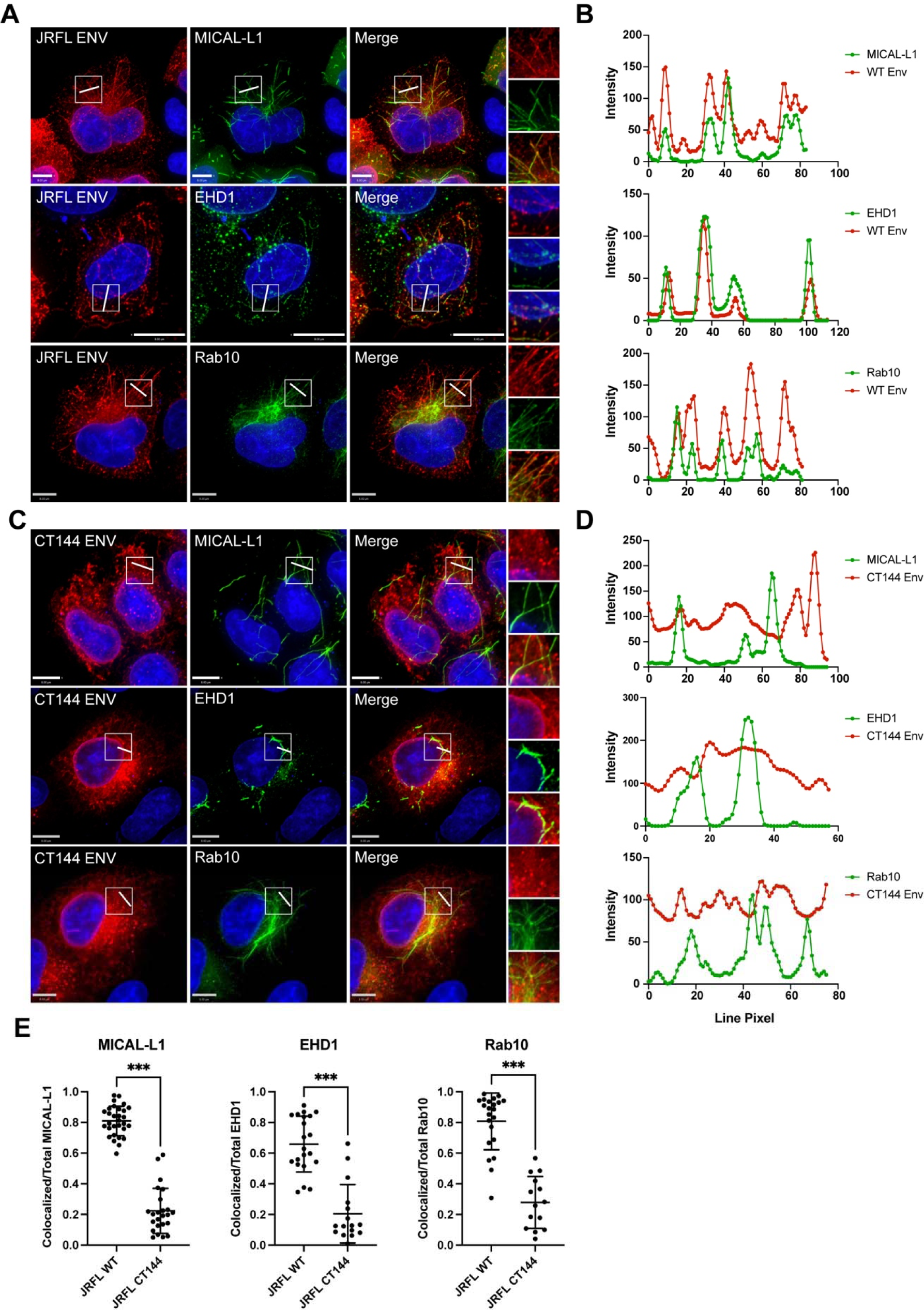
Env colocalization with the TRE requires an intact CT. (A) Env with a full-length CT was examined in cells together with staining for MICAL-L1, EHD1, and Rab10. A region of interest was selected for measuring intensity plots (boxed region and rightmost panels). (B) Linear intensity profiles were measured across tubules marked by TRE proteins, and compared with intensity of Env signal from (A). Note co-occurrence of intensity peaks of red and green pixels. (C) CT144 Env was expressed, and the procedure for generating intensity plots across TRE tubules was repeated. (D) Intensity plots for TRE markers (green) and CT144 Env (red), showing lack of concurrence of intensity peaks. (E) Manders’ correlation coefficient was measured as colocalized pixels/total pixels of each TRE marker from cells in (A) and (C) and is reported as mean ± SD. Significance was assessed using unpaired T-test. ***, P<0.001.

### FIP1C is present on the TRE and colocalizes with EHD1

FIP1C has previously been implicated as a trafficking adaptor involved in HIV-1 Env incorporation. We next asked if FIP1C is a component of the TRE. We utilized similar staining techniques to examine FIP1C distribution and colocalization with TRE markers. FIP1C has previously been noted to be enriched in the perinuclear ERC in HeLa cells along with a punctate cytoplasmic component.^19,30^ FIP1C was present in a markedly tubular distribution extending from the perinuclear region in a subset of cells examined following fixation methods preserving tubular endosomes (Figure 3A). Env and FIP1C colocalized on tubular membranes (Figure 3B). To determine if FIP1C is present on the TRE, we next examined colocalization with TRE markers. FIP1C appeared to colocalize strongly with EHD1 in these cells, showing a punctate distribution along the tubules marked by EHD1 (Figure 3C top, and quantitation, Figure 3D and 3E). Colocalization was not as prominent with MICAL-L1 (Figure 3C bottom, 3D, 3E). However, in these experiments FIP1C and MICAL-L1 tubules appeared to be continuous, with FIP1C somewhat more centrally located and MICAL-L1 more distal along the same tubules (Figure 3C, bottom). Together, these results indicate that FIP1C is a TRE component that colocalizes most profoundly with EHD1.

**Figure 3.**
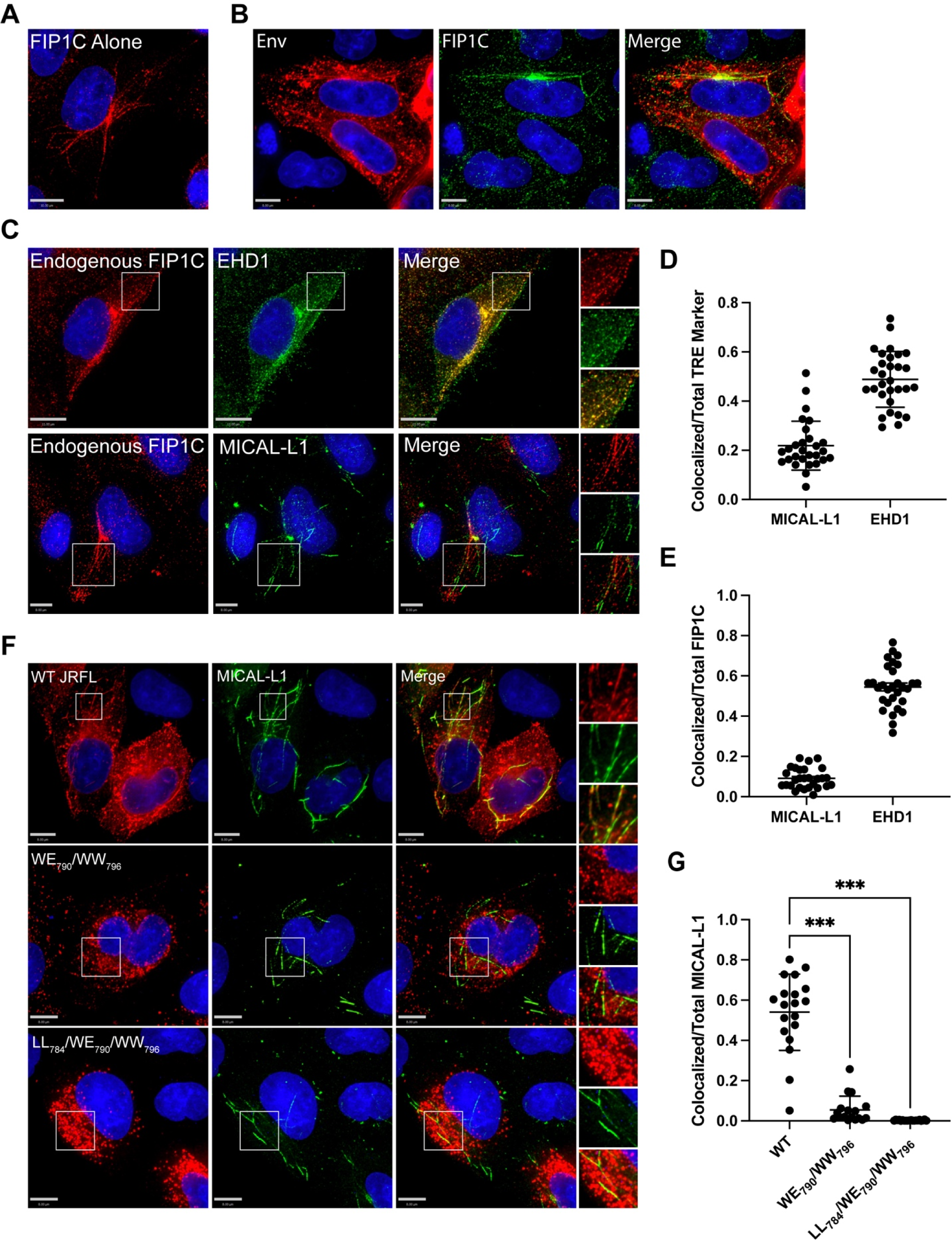
FIP1C is present on tubular endosomes and colocalizes with EHD1. (A) HeLa cells were stained for endogenous FIP1C. A subset of cells examined showed a clear tubular pattern of endogenous signal as shown. (B) FIP1C and Env colocalize on tubular endosomes. HeLa cells expressing JR-FL Env were fixed as before and stained for Env and FIP1C. Cells were fixed and stained as described previously. (C) Colocalization of FIP1C with TRE markers. Top Images: Striking colocalization was noted between endogenous FIP1C (red) and EHD1 (green) along tubular structures and in punctate form. Insets on right shown for clarity. Bottom images: endogenous FIP1C was found along tubules contiguous with MICAL-L1, with a lower degree of colocalization than that seen with EHD1. (D) Colocalization of FIP1C with EHD1 or MICAL-L1 as shown in (B) was assessed using Manders’ colocalization coefficient in multiple images, and shown as the proportion of colocalized pixels over total MICAL-L1 or EHD1 pixels, mean ±SD. (E) Analysis as in (D), but shown as colocalized pixels over total FIP1C pixels, mean ±SD. (F) Analysis of colocalization was performed with WT Env and MICAL-L1 (top) compared with two Env mutants bearing mutations in LLP3 region, WE_790_/WW_796_ (middle) and LL_784_/WE_790_/WW_796_^13^. (G) Manders’ correlation coefficients from experiment shown in (F) are plotted. Significance assessed using one way ANOVA with false discovery rate correction using the two stage set up method of Benjamini, Krieger, and Yekutieli^46^ included in Prism software (Graphpad). *** P<0.001.

### Tryptophan-based motifs in LLP3 regulate trafficking to TRE membranes

The N-terminal portion of the LLP3 region has been shown to play a role in Env incorporation in studies from several laboratories.^14–16^ We recently utilized a truncated FIP1C protein as an intervention to map motifs within the CT required for ERC localization. This work led to the identification of two tryptophan-based motifs in the N-terminal portion of LLP3 (WE_790_ and WW_796_) important for ERC localization, and disruption of both of these motifs in the proviral context led to significant defects in Env incorporation and viral replication.^13^ In order to determine if these motifs are important for TRE localization, we expressed two constructs bearing double tryptophan substitutions (WE_790_AA/WW_796_AA or LL_784_AA/WE_790_AA/WW_796_AA,^13^) in HeLa cells and examined the distribution of Env together with MICAL-L1. As described previously, WT Env was strongly colocalized with the TRE (Figure 3F, top panels). In contrast, both Env mutants failed to colocalize with the TRE (Figure 3F, middle and bottom panels). Using Manders’ correlation coefficient to measure overlapped pixels as a proportion of total MICAL-L1 signal, 54% of total MICAL-L1 signal was colocalized with WT Env, while only 5% and 0.2% overlap was observed for WE_790_AA/WW_796_AA or LL_784_AA/WE_790_AA/WW_796_AA, respectively (Figure 3G). Note that despite identical fixation techniques, the two mutant Envs were not observed in structures resembling tubular membranes. These results indicate that CT-dependent trafficking of Env to the TRE can be mapped to determinants in the N-terminal portion of LLP3.

### Disruption of the TRE leads to defects in Env incorporation in relevant (nonpermissive) cell types

Having established that HIV-1 Env traffics to the TRE in a CT-dependent manner, we next asked if perturbation of the TRE would inhibit Env incorporation. We chose to focus on depletion of MICAL-L1, a major scaffold protein and TRE component, and EHD1, an ATPase required for the scission of TRE membranes that enables vesicular delivery of cargo to the PM. We then evaluated the effects of TRE disruption on Env incorporation in multiple cell types, including those permissive for incorporation of CT-deleted Env (293T, MT-4), semipermissive HeLa cells, and three nonpermissive cell types (H9, CEM, Jurkat). After first establishing knockdown of MICAL-L1 or EHD1, cells were either transfected (293T, HeLa) or infected with VSV-G-pseudotyped NL4-3 virus (MT-4, H9, CEM, Jurkat). The production of Env was evaluated in cell lysates, and Env incorporation in released viral particles evaluated by pelleting of virus from supernatants. Depletion of MICAL-L1 or EHD1 was achieved in all cell lines evaluated (Figure 4A and 4B, MICAL-L1, and S1A and B, EHD1; compare scrambled vs. shRNA1 and shRNA2 or pooled lanes). No significant differences in Env production as marked by gp160 in cell lysates were observed (Figure 4A and 4B, cell lanes, MICAL-L1, and Figure S1A and S1B, EHD1). Incorporation of Env into particles was then assessed by comparing the amount of viral Env in scrambled vs target shRNA lanes, as represented by gp41 bands from virus particles (Figure 4A and 4B, S1A and S1B, asterisks). Despite a profound depletion of MICAL-L1 or EHD1, no reduction in Env incorporation into particles was seen for 293T, MT-4, or HeLa cells (Figure 4A and S1A, viral gp41 lanes). Remarkably, however, depletion of MICAL-L1 or EHD1 in the nonpermissive H9 T cell line caused a marked loss of Env incorporation (Figure 4B and S1B, H9, viral gp41 lanes). Loss of Env incorporation was also observed in MICAL-L1 or EHD1-depleted CEM cells (Figure 4B and S1B, CEM, viral gp41 lanes). Jurkat cells demonstrated a similar, though less profound, depletion of Env in released particles (Figure 4B and S1B, Jurkat, viral gp41 lanes). Quantitation of gp41/p24 ratios in released particles from permissive or semipermissive cells is shown in Figure 4C and S1C, and from nonpermissive cells in Figure 4D and S1D. As in the example blots shown, repeated experiments established that depletion of either MICAL-L1 or EHD1 led to loss of Env from particles released from H9, Jurkat, and CEM cells, with the most profound loss observed from H9 cells, while there was no loss of Env from HeLa, MT4, or 293T upon depletion of MICAL-L1 or EHD1. In summary of these experiments, disruption of TRE function by depletion of MICAL-L1 or EHD1 in permissive or semipermissive cells had no significant effect on Env incorporation, while disruption of TRE function in nonpermissive cells introduced a defect in Env incorporation, consistent with a model in which the TRE pathway is the dominant pathway for Env trafficking and incorporation in the most relevant cell types (nonpermissive cells).

**Figure 4.**
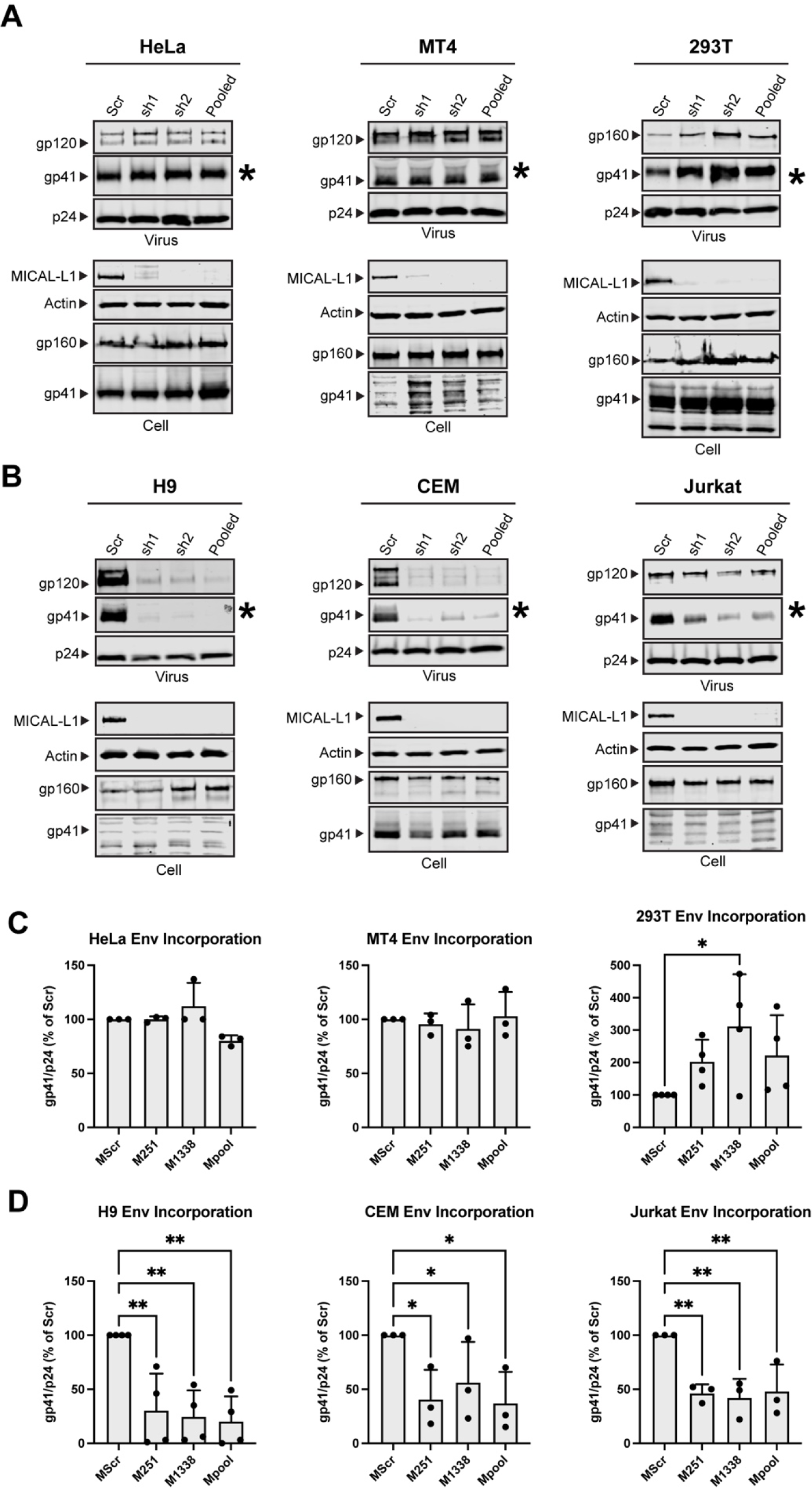
Depletion of MICAL-L1 reduces Env incorporation in a cell type-specific manner. Viral and cellular protein content was examined following shRNA-mediated knockdown of MICAL-L1 or treatment with scrambled (Scr) shRNA. (A) Semipermissive (HeLa) cells and permissive cells MT-4, 293T displayed no noticeable defect in Env incorporation despite high efficacy of knockdown. Asterisk denotes viral gp41 lane as indicator of Env incorporation into particles. (B) Nonpermissive cell types H9, CEM, and Jurkat cells demonstrate reduced particle incorporation of Env (gp41, asterisks) following MICAL-L1 depletion. (C) Quantitation of Env/p24 ratio, using Scr lanes as 100%, from repeated experiments; permissive/semipermissive cell panel. (D) Quantitation of Env/p24 ratio, using Scr lanes as 100%, from repeated experiments; nonpermissive cell panel. See also Figure S1 related to EHD1 depletion. Significance assessed using one way ANOVA with false discovery rate correction^46^ included in Prism software (Graphpad). * P<0.05, ** P<0.01.

### Cell Surface Levels of Env are unaffected by MICAL-L1, EHD1 depletion

Findings shown above indicate that Env incorporation into particles in nonpermissive T cell lines is dependent upon an intact TRE. We hypothesized that depletion of MICAL-L1 or EHD1 would reduce Env recycling to the PM, and therefore may reduce total cell surface envelope in these cell types. Cells were infected with VSV-G-pseudotyped HIV-1 (H9, MT-4, CEM, Jurkat) or transfected with NL4-3 proviral DNA (HeLa, 293T) following control shRNA or active shRNA depletion, and then assayed for cell surface Env by flow cytometry. Surprisingly, knockdown of either EHD1 or MICAL-L1 had no discernible effect on cell surface Env in any of the cell lines tested (Figure 5A). These surprising results were repeated a minimum of three times for each cell line. As discussed further below, this suggests that Env recycling from the TRE represents a small subset of total Env reaching the cell surface, and this Env subset is likely rapidly removed from the surface through particle budding.

**Figure 5.**
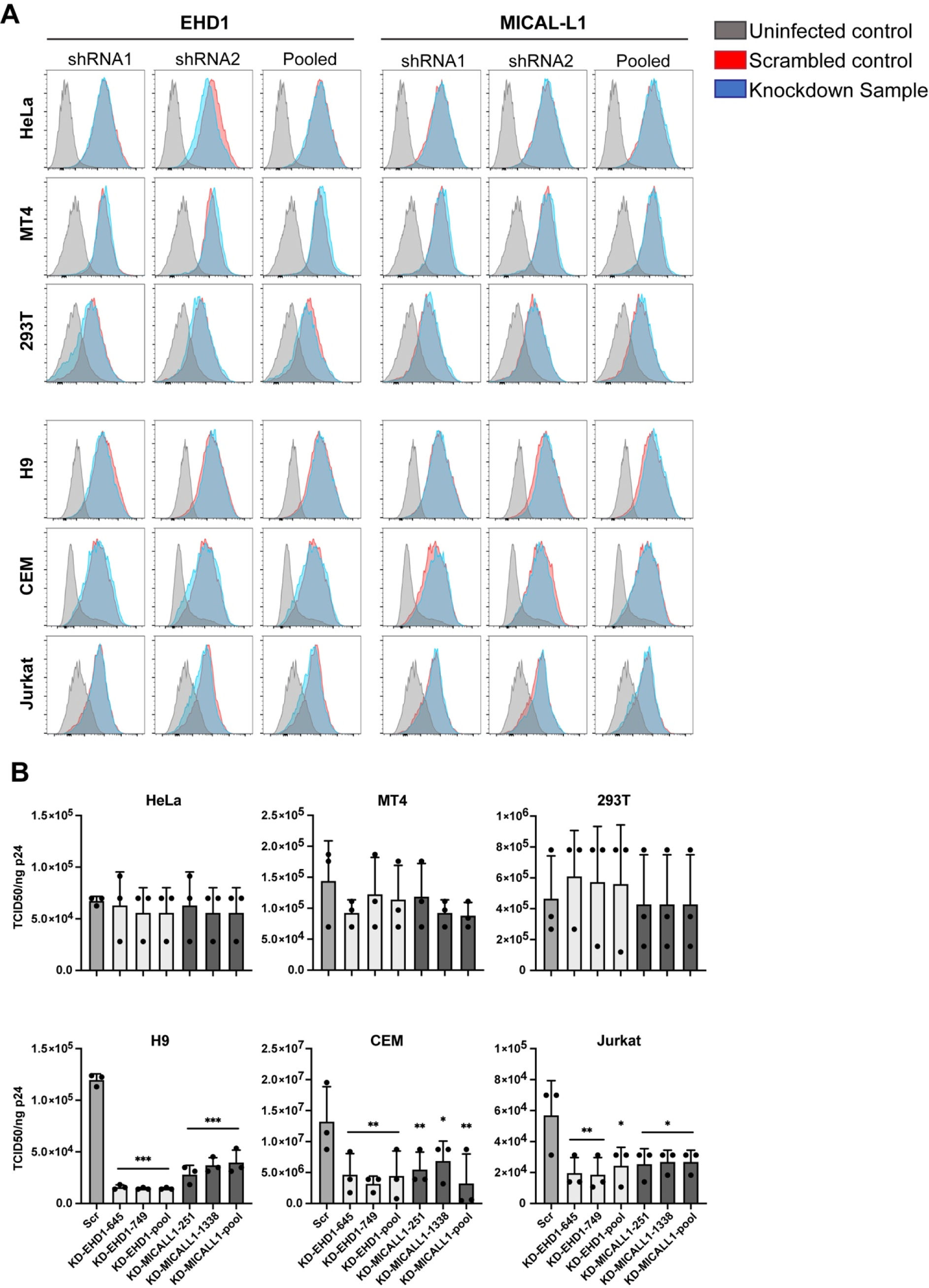
Knockdown of EHD1 or MICAL-L1 does not reduce cell surface Env but diminishes specific particle infectivity. (A) Knockdown or Scr cells were infected or transfected as described in the methods for Figure 5 and stained for cell surface Env with APC-conjugated 2G12 antibody. Cells were then permeabilized and stained for p24 with KC-57 FITC to gate for infected cells. Neither EHD1 nor MICAL-L1 knockdown (blue) reduced cell surface Env relative to Scr control (red) in any of the cell types tested. Grey indicates uninfected control cells stained with 2G12 antibody. (B) Virus was produced from EHD1 or MICAL-L1 knockdown cells as in experiments shown in Figure 4 and evaluated for specific infectivity by TZM-bl assay. EHD1 knockdown is shown in light colored bars, and MICAL-L1 knockdown is shown in the darkest colored bars. Semi-permissive HeLa and permissive 293T and MT4 cells showed no significant difference in particle infectivity from Scr treated or knockdown cells. In nonpermissive H9, CEM, and Jurkat cells there was a significant reduction in infectivity of particles produced following MICAL-L1 or EHD1 depletion. Significance was assessed using ANOVA with false discovery rate correction as before. *, P<0.05; **, P<0.01; ***, P<0.001.

### Cell type-dependence on an intact TRE for single-round infectivity and viral replication

To further evaluate the effect of TRE disruption on HIV particles, we examined single-round infectivity of particles produced from permissive or nonpermissive cells following EHD1 or MICAL-L1 depletion by inoculation onto TZM-bl reporter cells. Consistent with the Env incorporation results, neither EHD1 nor MICAL-L1 knockdown had an effect on infectivity of virus produced from 293T, MT4, or HeLa cells (Figure 5B). In contrast, depletion of MICAL-L1 or EHD1 in H9, CEM, or Jurkat cells significantly reduced particle infectivity; resulting in a decrease to 13%, 31%, and 37% for EHD1 knockdown or 29%, 47%, and 48% of control for the MICAL-L1 knockdown in H9, CEM, and Jurkat cells respectively (Figure 5B). Interestingly, the greatest effects in these experiments were observed following EHD1 depletion in H9 cells, with an average of 8.03-fold decrease in specific infectivity.

### Disruption of TRE components disrupts spreading HIV-1 infection in a cell type-dependent manner

Multiple-round assays of viral replication in cell culture may reveal different phenotypes than single-round assays, as both cell-cell and cell-free infection can contribute to ongoing viral spread. We evaluated the role of an intact TRE in MT-4, H9, CEM, and Jurkat cells following depletion of MICAL-L1 or EHD1 vs. control shRNA-treated cells. Cells were infected with VSV-G-pseudotyped NL4-3 virus, and production of virus monitored over time by measurement of p24 release. Depletion of either MICAL-L1 or EHD1 had no significant impact on viral replication in MT4 cells (Figure 6A and 6B). Remarkably, depletion of MICAL-L1 or EHD1 severely impaired viral replication in H9 cells (Figure 6C, 6D). Replication in CEM cells following depletion of MICAL-L1 or EHD1 also indicated a lower level of spreading infection upon TRE disruption (Figure 6E, 6F). A somewhat different pattern was observed with MICAL-L1 depletion in Jurkat cells, where an initial delay in viral propagation was observed followed by equivalent p24 production by the end of the growth curve sampling period (Figure 6G), while EHD1 depletion resulted in growth curves similar to those of CEM cells. The less profound depletion of Env from Jurkat or CEM cells upon depletion of MICAL-L1 (Figure 4) or EHD1 (Figure S1) likely explains the diminished magnitude of the effects seen in viral spread in Jurkat cells. These results suggest that the cell type-specific disruption of Env incorporation seen with disruption of TRE components leads to defects in both single-round infectivity and in the ability to replicate efficiently in a spreading infection.

**Figure 6.**
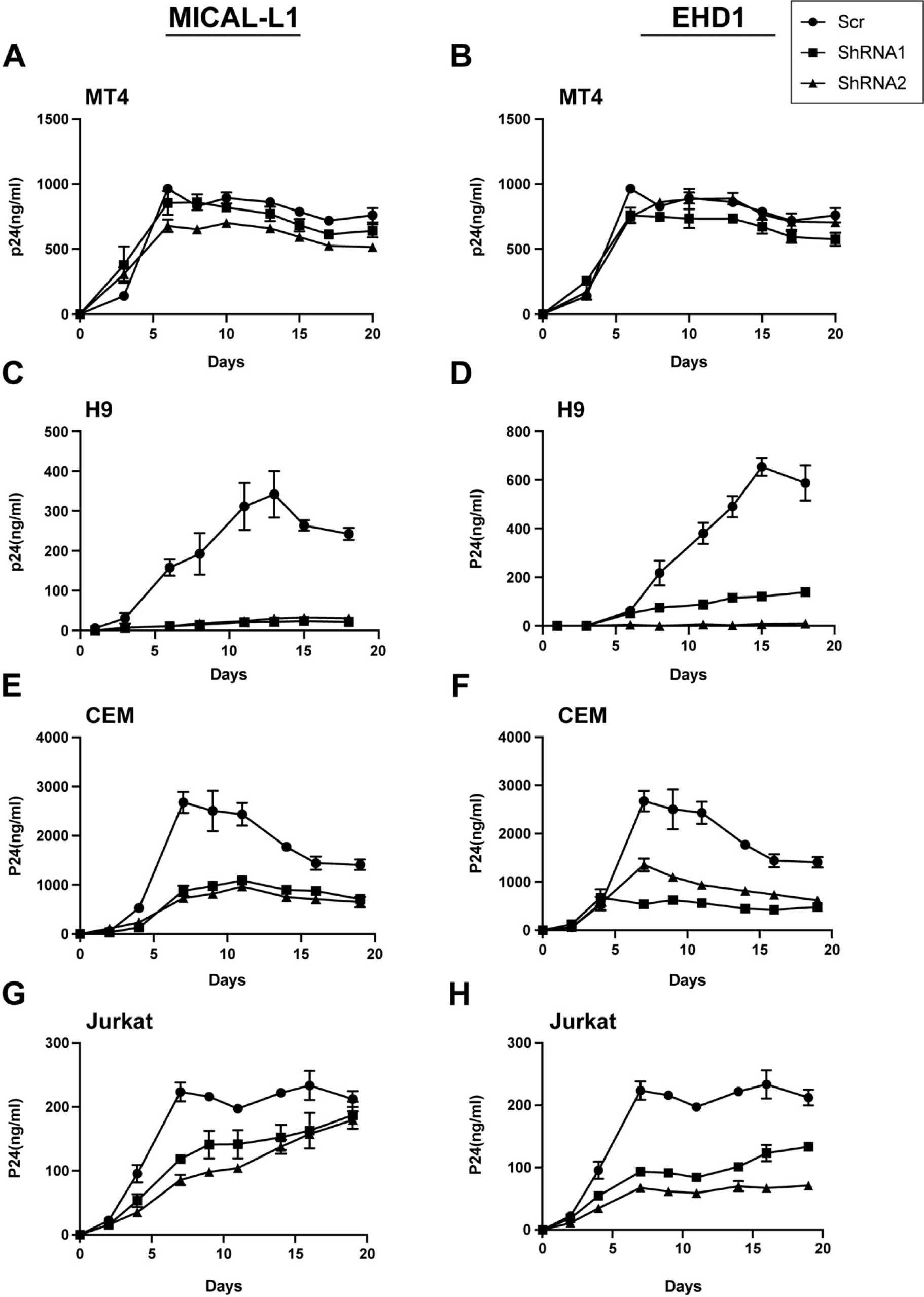
Depletion of MICAL-L1 or EHD1 results in replication defects in a cell type-specific manner. Cells were infected with NL4-3 and maintained for 3 weeks with intermittent sampling to assess virus release using p24 antigen assay. Growth curves following depletion of MICAL-L1 (left) or EHD1 (right) in permissive MT-4 cells (A,B), nonpermissive H9 cells (C,D), nonpermissive CEM cells (E,F) and nonpermissive Jurkat cells (G,H).

### Env trafficking to the TRE in primary monocyte-derived macrophages (MDMs)

The importance of the TRE in regulating Env incorporation in nonpermissive cells should reflect relevance to primary cells, as Env incorporation in primary T cells and macrophages requires an intact CT.^17^ To directly address this, we first asked if Env is found in the TRE within MDMs. MDMs were infected with the macrophage-tropic BaL strain of HIV, and fixed for immunofluorescence analysis for Env and markers of the TRE. MICAL-L1 was observed in tubular structures throughout the periphery of the MDMs, in a somewhat less organized pattern than that seen in HeLa cells (Figure 7A). Colocalization of Env was apparent in MICAL-L1 tubules (Figure 7A, inset panels). We next sought to deplete MICAL-L1 or EHD1 in primary MDMs. To do this, we treated MDMs with specific siRNAs or a scrambled siRNA control. A significant but not complete knockdown was achieved for both TRE components in these primary cells (Figure 7B, MICAL-L1, EHD1, siRNA lanes). Importantly, depletion of MICAL-L1 or EHD1 in primary MDMs reduced incorporation of gp41 and gp120 (Figure 7B, virus, gp41 and gp120 blots). This experiment was performed in MDMs from four different donors. Although there was some variability in gp41/p24 ratios between donors/experiments, a significant effect of MICAL-L1 or EHD1 depletion on Env incorporation into released particles was observed (Figure 7C). These results indicate that disruption of the TRE inhibits trafficking and particle incorporation in primary MDMs. When taken together with other results presented here, we conclude that Env trafficking and particle incorporation via the TRE is an important determinant of Env incorporation in the most relevant cell types for HIV replication.

**Figure 7.**
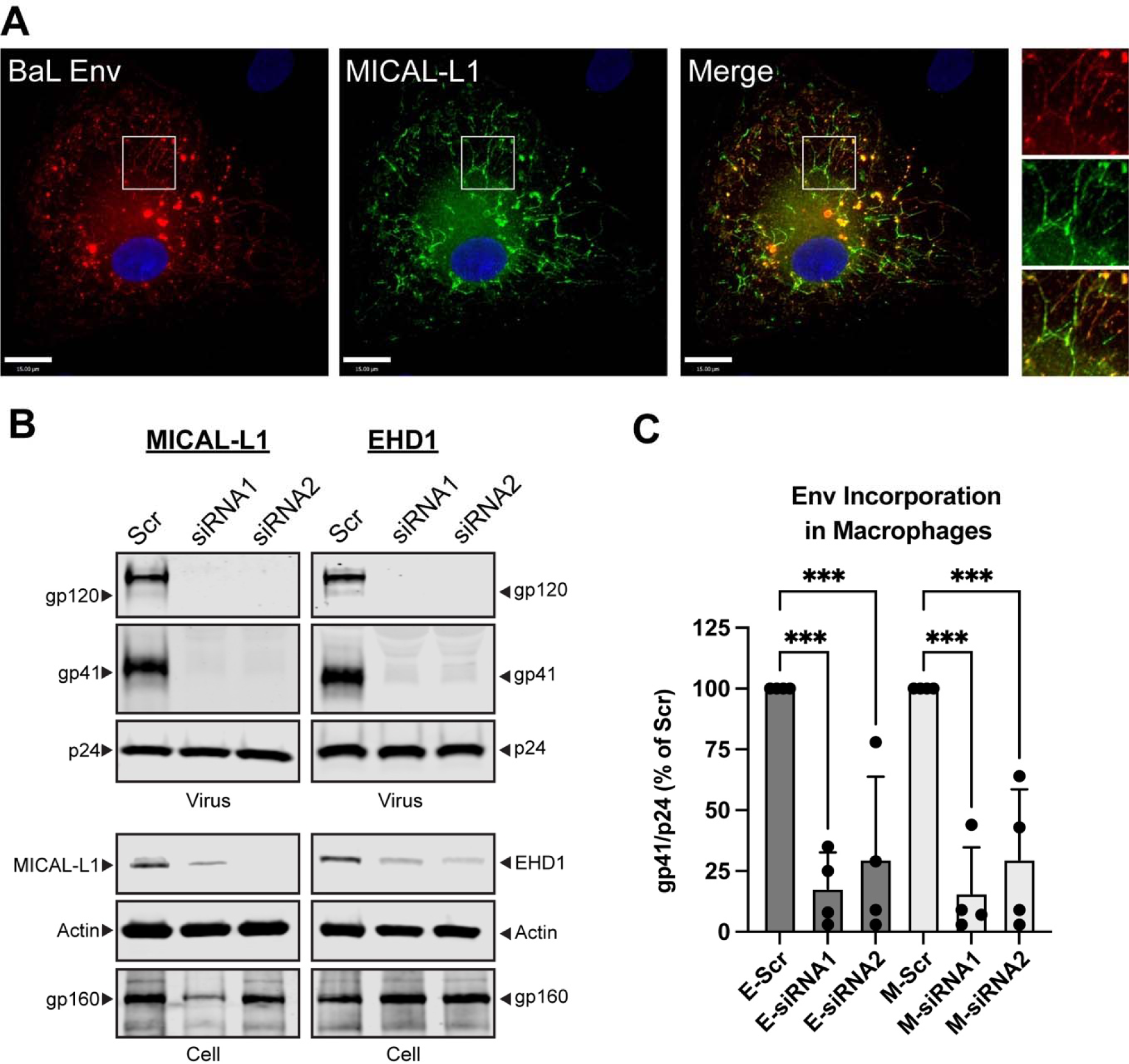
Env trafficking to the TRE regulates particle incorporation in primary MDMs. (A) MDMs were prepared from human donor monocytes and infected with HIV-1_BAL_. Cells were stained for HIV-1 Env and MICAL-L1 on day 5 post-infection. Region of interest shown as inset on right demonstrates colocalization of Env with this TRE marker. (B) siRNA-mediated depletion of MICAL-L1 or EHD1 was carried out in MDMs prior to infection with HIV-1_BAL_, using two distinct siRNAs and compared with Scr siRNA. Cell lysates and particles were harvested on day 7 following infection and examined by Western blot. Note depletion of gp41 and gp120 bands in siRNA lanes. (C) Quantitation of Env incorporation as assessed by gp41/p24 ratio in four separate experiments employing depletion of MICAL-L1 (M-siRNA) or EHD1 (E-siRNA). Significance addressed using one-way ANOVA as before. ***, P<0.001.

## DISCUSSION

The mechanism by which HIV-1 Env is incorporated into viral particles remains incompletely defined. Multiple models for Env incorporation have been proposed (reviewed in ^2,18^). Previous work from our laboratory and others has implicated a role for host cell trafficking pathways in regulating Env incorporation.^3,5,7,10,19,20,31^ It remains counterintuitive that HIV-1 Env is rapidly endocytosed to internal membranes where it may be shunted to the lysosome for degradation, delivered to the TGN through Golgi retrieval pathways, or recycled to the PM through the ERC, rather than simply remaining at the PM for subsequent particle assembly. Defining the precise trafficking pathways utilized by Env is important in order to determine whether specific host factors regulate the appearance of Env at membrane microdomains where interactions with the Gag lattice occur and infectious particles are generated. Furthermore, understanding Env trafficking may provide insights into immune evasion by HIV-infected cells, with implications for HIV cure and HIV vaccine development efforts.

Here we identify for the first time the interaction of HIV-1 Env with the TRE, a subcomponent of the cellular recycling apparatus that is responsible for sorting of receptors such as major histocompatibility complex class I (MHC-I) back to the PM following endocytosis. In live cells, Env at the PM rapidly entered tubules enriched in PIP2, and fixed cell studies confirmed that these tubules were membranes marked by the classical TRE components Rab10, MICAL-L1, and EHD1. TRE localization of Env required an intact CT, as truncated Env or Env with mutations of tryptophan-containing motifs in the LLP3 region failed to reach the TRE. The biological significance of this finding was confirmed by depletion of either MICAL-L1 or EHD1, which led to a significant reduction in Env incorporation and particle infectivity, and reduced replication in a spreading infection assay. Importantly, the effect of disruption of TRE components on Env incorporation was limited to specific cell types that have previously been described as “nonpermissive” for the incorporation of Env lacking an intact CT, thus linking the TRE to cell type-dependence of Env incorporation.

Murakami and Freed originally described the differences between cell types that require an intact CT for the efficient incorporation of Env into particles (nonpermissive cells) and cell types that allow incorporation of CT-deleted Env (permissive cells).^17^ It is important to note that replication of HIV-1 in the cell types most relevant to HIV-1 biology and pathogenesis, CD4+ T cells and macrophages, requires an intact CT (nonpermissive). These results suggest that host cell-specific factors that interact with the CT regulate Env incorporation. Evidence presented in this work establishes the TRE as an important determinant of cell type-dependent Env incorporation. This provides evidence for a model in which recycling to the PM determines particle incorporation of Env in the most relevant cell types. According to this model, endocytosed Env reaches the TRE, where sorting and interaction with specific TRE components determines subsequent scission from the TRE and recycling to the site of particle assembly for interaction with the developing Gag lattice. Differences in expression of essential components of the TRE required for interaction with the CT and recycling to the PM may therefore define differences between permissive and nonpermissive cells, and will require further dissection.

Depletion of MICAL-L1 or EHD1 led to significant reductions in Env particle incorporation, yet had no effect on the total cell surface Env in infected cells. This striking result suggest that it is only a small subfraction of the total cellular Env that is sorted back to the PM through TRE-dependent pathways. In our experiments using infection or expression of intact provirus, this fraction of Env at the PM would have been further depleted through particle incorporation and budding. It is well known that incorporation of Env into HIV particles is inefficient, resulting in only 7-14 trimers per virion.^32–34^ The relatively limited delivery of Env trimers from the TRE to the developing Gag lattice may determine this low level of incorporation, while the bulk of cell surface Env is somehow excluded from virion incorporation. It will be important in the future to define which components of the TRE represent the limiting factors in Env recycling and particle incorporation, allowing pursuit of interventions that either would inhibit or enhance delivery of Env to the site of assembly.

FIP1C is a cellular recycling adaptor previously implicated in Env trafficking and particle incorporation.^14,19,20^ A recent report found that FIP1C knockout did not limit HIV replication in CD4+ T cells, and questioned the relevance of this adaptor in explaining cell type-dependent incorporation of Env in primary cells.^21^ We note, however, that this report confirmed a significant effect of FIP1C on Env incorporation in several T cell lines including H9 cells. While FIP1C expression levels in nonpermissive vs. permissive cells do not explain differences in cell type-dependent incorporation of Env, the effects of FIP1C depletion on Env incorporation seen in cells such as H9, and the trapping of Env in the ERC upon overexpression of truncated FIP1C, established a connection between host cell recycling pathways and Env incorporation. The finding that FIP1C is also a TRE component and strongly colocalizes with EHD1 is intriguing, one that we suggest links this prior work to our current findings. We hypothesize that FIP1C is an adaptor that is involved in the sorting of an additional TRE-associated recycling factor that may itself interact with the CT. Notably, EHD1 and the related ATPase EHD3 have been documented to directly interact with Rab11-FIP2 (FIP2), and depletion of EHD1 alters FIP2 subcellular localization.^35^ FIP2 contains three NPF motifs, and binding to the second NPF motif of FIP2 is essential for EHD1 or EHD3 binding. While FIP1C has not been reported to directly bind to EHD1, the striking colocalization with EHD1 in this study suggests that it may associate with EHD1 either through a direct interaction or through heterodimerization with FIP2.^36^

EHD1 is an ATPase involved in scission of membranes, facilitating vesicular delivery of proteins such as transferrin receptor, β1 integrin, the glucose transporter GLUT4, epidermal growth factor receptor, and MHC1 from the TRE to the PM.^22,37–40^ EHD1 is recruited to the TRE through interactions with NPF motifs present on MICAL-L1 and syndapin2, a MICAL-L1 binding partner and F-BAR domain protein. The involvement of EHD1 in facilitating delivery of a subpopulation of cellular HIV-1 Env to the PM for incorporation into particles raises additional questions regarding how Gag and Env interact within the cell. PIP2- and Env-enriched vesicles released from the TRE and transported to the PM could constitute natural binding sites for Gag through the interaction of the basic patch on the matrix domain of Gag with PIP2-enriched membranes.^41^ Interactions of Gag with PIP2-enriched membranes on the TRE prior to EHD1-mediated scission are also possible. It is of interest that the MICAL-L1- and EHD1-binding partner syndapin2 (also known as pacsin2) is itself incorporated into HIV-1 particles through an interaction with the p6 domain of Gag, and has been previously implicated in facilitating HIV spreading infection.^42^

In summary, we have shown that trafficking of HIV-1 Env to the TRE is CT-dependent, and is a defining factor in cell type-dependent incorporation of Env into particles. This work provides support for a model in which host recycling factors are crucial for particle incorporation, and opens up new avenues for investigating this step in the HIV-1 lifecycle.

## MATERIALS AND METHODS

### Cell Lines

HeLa, 293T, H9, Jurkat, and CEM cells were obtained from the American Type Culture Collection (ATCC; CCL-2, CRL-3216, HTB-176, TIB-152, and CCL-119). MT-4 cells were obtained from the NIH AIDS Reagent Program (ARP-120). TZM-bl cells were obtained through the NIH AIDS Reagent Program, Division of AIDS, NIAID, NIH, from John C. Kappes, Xiaoyun Wu, and Tranzyme, Inc. HeLa, TZM-bl, and 293T cells were maintained in Dulbecco’s modified Eagle Medium (DMEM) (Thermo Fisher Scientific) supplemented with 10% FBS (Fetal Bovine Serum), 2 mM L-glutamine, 100 IU penicillin, and 100 μg/mL streptomycin. H9, Jurkat, CEM, and MT-4 cells were maintained in RPMI 1640 (Roswell Park Memorial Institute) containing 10% FBS (Fetal Bovine Serum), 2 mM L-glutamine, 100 IU penicillin, and 100 μg/mL streptomycin. All cell lines were cultured from early passage frozen stocks from the original source and were documented to be mycoplasma negative.

### Primary MDM culture

Primary monocyte-derived macrophages (MDMs) were prepared as follows: peripheral blood mononuclear cells (PBMCs) were isolated from donors using Ficoll-Hypaque gradient centrifugation. PBMCs from the buffy coats were pooled and extensively washed with PBS to remove residual platelets. Monocytes were enriched by negative selection method using Miltenyi Pan monocyte isolation kit (Miltenyi Biotec Inc). Enriched monocytes were plated on poly-D-lysine coated 35mm Mattek dishes (MatTek) or poly-D-lysine coated 6 well plates (Corning) and cultured in RPMI 1640 media supplemented with 10% FBS (Bio-Techne), 100U/ml penicillin, 100ug/ml streptomycin and 2mM Glutamine. The cells were matured in the presence of 5 ng/ml GM-CSF (PeproTech) for 7 days to facilitate maturation into monocyte derived macrophages (MDMs). PBMCs were obtained from four distinct donors for this analysis.

### shRNA or siRNA-mediated knockdown

Knockdowns were performed in all cells except MDMs using lentiviral transduction of cells with shRNA. ShRNA plasmids were acquired from Sigma for EHD1 and MICAL-L1. To produce lentiviral particles, 293T cells were seeded at a density of one million cells per well of a 6-well plate overnight. The next day, cells were transfected with 0.5 μg pMD2.G, 0.5 μg psPAX2, and 1 μg of MICAL-L1 or EHD1 shRNA in pLKO.1 vector using Jetprime transfection reagent (Polyplus). Following transfection, 293T cells were incubated for 48 hours then harvested and clarified by centrifugation and filtration through 0.45 μm filter. 250,000 HeLa, 1 million 293T, or 3-4 million H9, Jurkat, MT4, or CEM were infected overnight in a 6-well plate in the presence of 0.5 μg/mL polybrene (Sigma-Aldrich). The following day, cells were incubated with selection media containing 1 μg/mL puromycin (H9, CEM, MT-4) or 2 μg/mL puromycin for Jurkat. The degree of knockdown of each target gene in each cell type was then assessed by Western blot analysis. For knockdown of MICAL-L1 or EHD1 in MDMs, siRNA-mediated depletion was performed with RNAiMAX reagents purchased from Sigma. Macrophages were serum starved for one hour prior to transfection. 25 nmol of siRNA was diluted in 100 uL of Optimem media (Thermo Fisher Scientific) while in a separate tube 20 uL of Lipofectamine RNAiMAX transfection reagent (Thermo Fisher Scientific) was mixed with another 100 uL of OptiMem media. The diluted siRNA and RNAiMAX stocks were mixed 1:1 and incubated for 5 minutes at room temperature followed by addition to the Macrophages in serum free media. After 6-8 hours, Macrophages were switched to RPMI 1640 media supplemented with 10% FBS, 100U/ml penicillin, 100ug/ml streptomycin and 2mM Glutamine.

### Viral infection for analysis of Env incorporation

4-5 million MICAL-L1 specific, EHD1 specific, or scrambled shRNA-treated T-cells were infected with 150 ng p24 of VSV-G-pseudotyped NL4-3 virus for 48 hours. HeLa and 293T scrambled shRNA-treated or knockdown cells were plated at a density of 250,000 cells per well of a 6 well plate and transfected with 0.5 μg of NL4-3 plasmid DNA overnight. Supernatants were precleared of cells and debris by centrifuging at 17,000 rcf for 5 minutes. Viruses were harvested from supernatants by pelleting through a 20% sucrose cushion at maximum speed (17,000 rcf) for 2 hours on a microcentrifuge. Viral pellets were then lysed in ice cold RIPA buffer with protease inhibitors. Cells were also lysed in RIPA buffer. Quantitation of virus stocks was performed by p24 ELISA as previously described.^43^ For infection of MDMs, a stock of HIV-1_BAL_ prepared from human PBMCs stimulated with PHA and IL-2 was prepared as previously described,^44^ and MDMs were infected at an MOI of 0.75 as measured on TZM-bl reporter cells. Imaging was performed 5 days post-infection. Harvesting of MDMs for analysis of cell lysates and viral pellets was performed on day 7 post-infection.

### Western blotting for viral and cellular proteins

Viruses were pelleted through a 20% sucrose cushion and resuspended in RIPA buffer, and cells were lysed with RIPA buffer. Viral supernatants were normalized for p24 by ELISA, and cell lysates were normalized for total protein by DC protein assay. Supernatants and lysates were analyzed by 10% bis-tris SDS-PAGE and transferred to nitrocellulose membranes. Membranes were blocked with Intercept blocking buffer (LI-Cor Biosciences) for 30 minutes followed by antibody staining. All antibodies were diluted in Intercept blocking buffer with 0.15% Tween-20 at the following dilutions: gp41 was detected with human 2F5 (Polymun) (1:1000); gp120/gp160 were detected with human antibody 2G12 (Polymun) (1:1000); actin was detected with mouse anti-actin (Thermo Fisher Scientific, MA5-11869) (1:3000); p24 was detected with mouse CA183 (1:1000); MICAL-L1 was detected with mouse anti-MICAL-L1 (Novusbio, H00085377-B01P) (1:300); EHD1 was detected with rabbit anti-EHD1 (Sigma-Aldrich, 067747) (1:500). Primary antibodies were detected with LICOR IRDyes (LI-Cor Biosciences) against the appropriate species diluted 1:10,000 in blocking buffer with 0.15% Tween-20.

### Infectivity measurement and replication assays

Infectivity of cell culture supernatants was measured using TZM-bl indicator cells following p24 normalization as previously described.^45^ For assessment of multi-round replication, 1 million MICAL-L1 KD, EHD1 KD, or scrambled shRNA-transduced T-cells were infected with 50 ng of VSV-G-pseudotyped NL4-3 virus overnight. The next day, the cells were washed and plated in 12-well dishes in 2 mL of RPMI containing 10% FBS supplemented with 1ug/ml puromycin and antibiotics. Every two to three days, 200 μL of media was removed and replaced with fresh media. Sampled supernatants were analyzed by p24 ELISA.

### Immunofluorescence microscopy

For live cell experiments, HeLa cells were plated in 35mm^2^ poly–d-lysine-treated dishes (MatTek) overnight and then were then transfected with plcδ-PH-GFP and pcDNA5/TO JR-FLoptgp160-FAP constructs using jetPRIME (Polyplus). 20 hours later, live cell imaging was conducted in 5% CO_2_ at 37°C chamber on a Deltavision Elite live cell deconvolution imaging system using the TIRF imaging configuration. FAP reagent αRED-np1 (Spectragenetics) was directly added to the supernatant as images were acquired, with images taken at a rate one every 15 seconds over a period of 20 minutes. For imaging of fixed samples, cells were washed with warm PBS then fixed for fifteen minutes with 4% paraformaldehyde in PBS prewarmed to 37°C. Following fixation, samples were permeabilized by briefly incubating with 0.05% Triton X-100 (Sigma Aldrich), then blocked with Dako protein block (Agilent Technologies) for 30 minutes. Blocking solution was washed out with PBS containing 0.05% Triton X-100 and samples were stained with primary antibody diluted in Dako antibody diluent (Agilent Technologies). Primary antibodies and dilutions used were as follows: 2G12 for Env (1:1000); EHD1 (1:500); MICAL-L1 (1:300). Live imaging was performed using FAP-Env and GFP-PLCδ1-PH expressed for 24 hours. FAP-Env was incubated with non-membrane permeable fluorogenic peptide reagent for 5 minutes prior to wash out and imaging. Imaging was performed with Deltavision Elite or Deltavision OMX microscopes (Leica Microsystems, Wetzlar, Germany) with a 60X lens (NA: 1.42) and images were processed with Volocity software (Quorum Technologies, Inc).

### Flow Cytometry

Cells were infected with VSV-G-pseudotyped NL4-3 overnight followed by wash with PBS and incubation at 37°C for 24-48 hours. Cells were then collected into microcentrifuge tubes, stained with zombie violet diluted 1:500 in PBS for live dead discrimination for 30 minutes at room temperature, washed twice with PBS, and fixed with 4% paraformaldehyde for 10 minutes. Blocking was performed with Dako protein block supplemented with 1:100 dilution of 1 mg/mL non-specific human IgG for 30 minutes. Block was washed out with PBS, and cells were incubated with 2G12 directly conjugated to APC (Abcam, ab201807) diluted 1:100 from 1 mg/mL in Dako antibody diluent for 2 hours to stain for cell surface Env. Cells were washed twice and permeabilized with 0.2% Triton X-100 for five minutes and washed again. Cells were then stained for Gag with KC57-FITC (Beckman Coulter, 6604665) for 2 hours, washed twice with PBS, and were resuspended in an appropriate amount of MACS buffer (Miltenyi Biotec) supplemented with BSA (Miltenyi Biotec). Samples were analyzed using BD FACS Canto II (BD Biosciences) and data was processed with FlowJo software (BD Biosciences).

### Statistical Analysis

Manders’ colocalization coefficient was performed using the Volocity software package (Quorum Technologies, Inc). Comparisons between colocalization values from multiple cells were plotted as mean ±SD. Significance was determined using unpaired t-test for colocalization differences in Figure 2, employing Graphpad Prism for these calculations. In all other comparisons where multiple comparisons were performed, one way ANOVA with false discovery rate correction^46^ was performed using Graphpad Prism.

## AUTHOR CONTRIBUTIONS

Conceptualization, G.L. and P.S., Methodology, G.L., P.S., Investigation, G.L., L.D., and K.C., Resources, G.L. and K.C., Writing-original draft G.L.; Writing-Reviewing and Editing, P.S., L.D., K.C., and G.L., Funding Acquisition, P.S., Supervision, P.S.

## ACKNOWLEDGMENTS

This work was supported by R01AI150486.

The funders had no role in study design, data collection and interpretation, or the decision to submit the work for publication. Flow cytometry was performed using the Cincinnati Children’s Hospital Medical Center (CCHMC) Flow Cytometry Core.

## REFERENCES

1. Bernstein, H.B., Tucker, S.P., Hunter, E., Schutzbach, J.S., and Compans, R.W. (1994). Human immunodeficiency virus type 1 envelope glycoprotein is modified by O-linked oligosaccharides. Journal of Virology 68, 463–468. doi:10.1128/jvi.68.1.463-468.1994.

2. Checkley, M.A., Luttge, B.G., and Freed, E.O. (2011). HIV-1 Envelope Glycoprotein Biosynthesis, Trafficking, and Incorporation. Journal of Molecular Biology 410, 582–608. 10.1016/j.jmb.2011.04.042.

3. Willey, R.L., Bonifacino, J.S., Potts, B.J., Martin, M.A., and Klausner, R.D. (1988). Biosynthesis, cleavage, and degradation of the human immunodeficiency virus 1 envelope glycoprotein gp160. Proceedings of the National Academy of Sciences 85, 9580–9584. doi:10.1073/pnas.85.24.9580.

4. Hallenberger, S., Bosch, V., Angliker, H., Shaw, E., Klenk, H.D., and Garten, W. (1992). Inhibition of furin-mediated cleavage activation of HIV-1 glycoprotein gp160. Nature 360, 358–361. 10.1038/360358a0.

5. Egan, M.A., Carruth, L.M., Rowell, J.F., Yu, X., and Siliciano, R.F. (1996). Human immunodeficiency virus type 1 envelope protein endocytosis mediated by a highly conserved intrinsic internalization signal in the cytoplasmic domain of gp41 is suppressed in the presence of the Pr55gag precursor protein. J Virol 70, 6547–6556. 10.1128/JVI.70.10.6547-6556.1996.

6. Ohno, H., Aguilar, R.C., Fournier, M.-C., Hennecke, S., Cosson, P., and Bonifacino, J.S. (1997). Interaction of Endocytic Signals from the HIV-1 Envelope Glycoprotein Complex with Members of the Adaptor Medium Chain Family. Virology 238, 305–315. 10.1006/viro.1997.8839.

7. Boge, M., Wyss, S., Bonifacino, J.S., and Thali, M. (1998). A Membrane-proximal Tyrosine-based Signal Mediates Internalization of the HIV-1 Envelope Glycoprotein via Interaction with the AP-2 Clathrin Adaptor *. Journal of Biological Chemistry 273, 15773–15778. 10.1074/jbc.273.25.15773.

8. Rowell, J.F., Stanhope, P.E., and Siliciano, R.F. (1995). Endocytosis of endogenously synthesized HIV-1 envelope protein. Mechanism and role in processing for association with class II MHC. The Journal of Immunology 155, 473–488.

9. Wyss, S., Berlioz-Torrent, C., Boge, M., Blot, G., Höning, S., Benarous, R., and Thali, M. (2001). The highly conserved C-terminal dileucine motif in the cytosolic domain of the human immunodeficiency virus type 1 envelope glycoprotein is critical for its association with the AP-1 clathrin adaptor [correction of adapter]. J Virol 75, 2982–2992. 10.1128/jvi.75.6.2982-2992.2001.

10. Byland, R., Vance, P.J., Hoxie, J.A., and Marsh, M. (2007). A conserved dileucine motif mediates clathrin and AP-2-dependent endocytosis of the HIV-1 envelope protein. Mol Biol Cell 18, 414–425. 10.1091/mbc.e06-06-0535.

11. Groppelli, E., Len, A.C., Granger, L.A., and Jolly, C. (2014). Retromer regulates HIV-1 envelope glycoprotein trafficking and incorporation into virions. PLoS Pathog 10, e1004518–e1004518. 10.1371/journal.ppat.1004518.

12. Bültmann, A., Muranyi, W., Seed, B., and Haas, J. (2001). Identification of two sequences in the cytoplasmic tail of the human immunodeficiency virus type 1 envelope glycoprotein that inhibit cell surface expression. Journal of virology 75, 5263–5276. 10.1128/JVI.75.11.5263-5276.2001.

13. Lerner, G., Ding, L., and Spearman, P. (2023). Tryptophan-based motifs in the LLP3 region of the HIV-1 envelope glycoprotein cytoplasmic tail direct trafficking to the endosomal recycling compartment and mediate particle incorporation. Journal of Virology 0, e00631–00623. 10.1128/jvi.00631-23.

14. Qi, M., Chu, H., Chen, X., Choi, J., Wen, X., Hammonds, J., Ding, L., Hunter, E., and Spearman, P. (2015). A tyrosine-based motif in the HIV-1 envelope glycoprotein tail mediates cell-type- and Rab11-FIP1C-dependent incorporation into virions. Proc Natl Acad Sci U S A 112, 7575–7580. 10.1073/pnas.1504174112.

15. Bhakta, S.J., Shang, L., Prince, J.L., Claiborne, D.T., and Hunter, E. (2011). Mutagenesis of tyrosine and di-leucine motifs in the HIV-1 envelope cytoplasmic domain results in a loss of Env-mediated fusion and infectivity. Retrovirology 8, 37. 10.1186/1742-4690-8-37.

16. Murakami, T., and Freed, E.O. (2000). Genetic evidence for an interaction between human immunodeficiency virus type 1 matrix and alpha-helix 2 of the gp41 cytoplasmic tail. J Virol 74, 3548–3554. 10.1128/jvi.74.8.3548-3554.2000.

17. Murakami, T., and Freed, E.O. (2000). The long cytoplasmic tail of gp41 is required in a cell type-dependent manner for HIV-1 envelope glycoprotein incorporation into virions. Proc Natl Acad Sci U S A 97, 343–348. 10.1073/pnas.97.1.343.

18. Anokhin, B., and Spearman, P. (2022). Viral and Host Factors Regulating HIV-1 Envelope Protein Trafficking and Particle Incorporation. Viruses 14. 10.3390/v14081729.

19. Qi, M., Williams, J.A., Chu, H., Chen, X., Wang, J.-J., Ding, L., Akhirome, E., Wen, X., Lapierre, L.A., Goldenring, J.R., and Spearman, P. (2013). Rab11-FIP1C and Rab14 Direct Plasma Membrane Sorting and Particle Incorporation of the HIV-1 Envelope Glycoprotein Complex. PLoS Pathog 9, e1003278. 10.1371/journal.ppat.1003278.

20. Kirschman, J., Qi, M., Ding, L., Hammonds, J., Dienger-Stambaugh, K., Wang, J.J., Lapierre, L.A., Goldenring, J.R., and Spearman, P. (2018). HIV-1 Envelope Glycoprotein Trafficking through the Endosomal Recycling Compartment Is Required for Particle Incorporation. J Virol 92, DOI: 10.1128/JVI.01893-01817. 10.1128/JVI.01893-17.

21. Fernandez-de Cespedes, M.V., Hoffman, H.K., Carter, H., Simons, L.M., Naing, L., Ablan, S.D., Scheiblin, D.A., Hultquist, J.F., van Engelenburg, S.B., and Freed, E.O. (2022). Rab11-FIP1C Is Dispensable for HIV-1 Replication in Primary CD4(+) T Cells, but Its Role Is Cell Type Dependent in Immortalized Human T-Cell Lines. J Virol 96, e0087622. 10.1128/jvi.00876-22.

22. Caplan, S., Naslavsky, N., Hartnell, L.M., Lodge, R., Polishchuk, R.S., Donaldson, J.G., and Bonifacino, J.S. (2002). A tubular EHD1-containing compartment involved in the recycling of major histocompatibility complex class I molecules to the plasma membrane. The EMBO Journal 21, 2557–2567. 10.1093/emboj/21.11.2557.

23. Farmer, T., Xie, S., Naslavsky, N., Stöckli, J., James, D.E., and Caplan, S. (2021). Defining the protein and lipid constituents of tubular recycling endosomes. Journal of Biological Chemistry 296, 100190. 10.1074/jbc.RA120.015992.

24. Jović, M., Kieken, F., Naslavsky, N., Sorgen, P.L., and Caplan, S. (2009). Eps15 Homology Domain 1-associated Tubules Contain Phosphatidylinositol-4-Phosphate and Phosphatidylinositol-(4,5)-Bisphosphate and Are Required for Efficient Recycling. Molecular Biology of the Cell 20, 2731–2743. 10.1091/mbc.e08-11-1102.

25. Etoh, K., and Fukuda, M. (2019). Rab10 regulates tubular endosome formation through KIF13A and KIF13B motors. J Cell Sci 132. 10.1242/jcs.226977.

26. Weaver, N., Hammonds, J., Ding, L., Lerner, G., Dienger-Stambaugh, K., and Spearman, P. (2023). KIF16B Mediates Anterograde Transport and Modulates Lysosomal Degradation of the HIV-1 Envelope Glycoprotein. Journal of Virology 97, e00255–00223. 10.1128/jvi.00255-23.

27. Boucrot, E., Ferreira, A.P., Almeida-Souza, L., Debard, S., Vallis, Y., Howard, G., Bertot, L., Sauvonnet, N., and McMahon, H.T. (2015). Endophilin marks and controls a clathrin-independent endocytic pathway. Nature 517, 460–465. 10.1038/nature14067.

28. Day, C.A., Baetz, N.W., Copeland, C.A., Kraft, L.J., Han, B., Tiwari, A., Drake, K.R., De Luca, H., Chinnapen, D.J., Davidson, M.W., et al. (2015). Microtubule motors power plasma membrane tubulation in clathrin-independent endocytosis. Traffic 16, 572–590. 10.1111/tra.12269.

29. Xie, S., Bahl, K., Reinecke, J.B., Hammond, G.R., Naslavsky, N., and Caplan, S. (2016). The endocytic recycling compartment maintains cargo segregation acquired upon exit from the sorting endosome. Mol Biol Cell 27, 108–126. 10.1091/mbc.E15-07-0514.

30. Peden, A.A., Schonteich, E., Chun, J., Junutula, J.R., Scheller, R.H., and Prekeris, R. (2004). The RCP-Rab11 complex regulates endocytic protein sorting. Mol Biol Cell 15, 3530–3541. 10.1091/mbc.e03-12-0918.

31. Hoffman, H.K., Aguilar, R.S., Clark, A.R., Groves, N.S., Pezeshkian, N., Bruns, M.M., and van Engelenburg, S.B. (2022). Endocytosed HIV-1 Envelope Glycoprotein Traffics to Rab14(+) Late Endosomes and Lysosomes to Regulate Surface Levels in T-Cell Lines. J Virol 96, e0076722. 10.1128/jvi.00767-22.

32. Chertova, E., Bess, J.W., Jr., Crise, B.J., Sowder, I.R., Schaden, T.M., Hilburn, J.M., Hoxie, J.A., Benveniste, R.E., Lifson, J.D., Henderson, L.E., and Arthur, L.O. (2002). Envelope glycoprotein incorporation, not shedding of surface envelope glycoprotein (gp120/SU), Is the primary determinant of SU content of purified human immunodeficiency virus type 1 and simian immunodeficiency virus. J Virol 76, 5315–5325. 10.1128/jvi.76.11.5315-5325.2002.

33. Zhu, P., Chertova, E., Bess, J., Jr., Lifson, J.D., Arthur, L.O., Liu, J., Taylor, K.A., and Roux, K.H. (2003). Electron tomography analysis of envelope glycoprotein trimers on HIV and simian immunodeficiency virus virions. Proc Natl Acad Sci U S A 100, 15812–15817. 10.1073/pnas.2634931100.

34. Zhu, P., Liu, J., Bess, J., Jr., Chertova, E., Lifson, J.D., Grise, H., Ofek, G.A., Taylor, K.A., and Roux, K.H. (2006). Distribution and three-dimensional structure of AIDS virus envelope spikes. Nature 441, 847–852. 10.1038/nature04817.

35. Naslavsky, N., Rahajeng, J., Sharma, M., Jovic, M., and Caplan, S. (2006). Interactions between EHD proteins and Rab11-FIP2: a role for EHD3 in early endosomal transport. Mol Biol Cell 17, 163–177. 10.1091/mbc.e05-05-0466.

36. Wallace, D.M., Lindsay, A.J., Hendrick, A.G., and McCaffrey, M.W. (2002). The novel Rab11-FIP/Rip/RCP family of proteins displays extensive homo- and hetero-interacting abilities. Biochem Biophys Res Commun 292, 909–915. 10.1006/bbrc.2002.6736.

37. Guilherme, A., Soriano, N.A., Furcinitti, P.S., and Czech, M.P. (2004). Role of EHD1 and EHBP1 in perinuclear sorting and insulin-regulated GLUT4 recycling in 3T3-L1 adipocytes. J Biol Chem 279, 40062–40075. 10.1074/jbc.M401918200.

38. Tom, E.C., Mushtaq, I., Mohapatra, B.C., Luan, H., Bhat, A.M., Zutshi, N., Chakraborty, S., Islam, N., Arya, P., Bielecki, T.A., et al. (2020). EHD1 and RUSC2 Control Basal Epidermal Growth Factor Receptor Cell Surface Expression and Recycling. Mol Cell Biol 40. 10.1128/MCB.00434-19.

39. Jovic, M., Naslavsky, N., Rapaport, D., Horowitz, M., and Caplan, S. (2007). EHD1 regulates beta1 integrin endosomal transport: effects on focal adhesions, cell spreading and migration. J Cell Sci 120, 802–814. 10.1242/jcs.03383.

40. Naslavsky, N., Boehm, M., Backlund, P.S., Jr., and Caplan, S. (2004). Rabenosyn-5 and EHD1 interact and sequentially regulate protein recycling to the plasma membrane. Mol Biol Cell 15, 2410–2422. 10.1091/mbc.e03-10-0733.

41. Saad, J.S., Miller, J., Tai, J., Kim, A., Ghanam, R.H., and Summers, M.F. (2006). Structural basis for targeting HIV-1 Gag proteins to the plasma membrane for virus assembly. Proc Natl Acad Sci U S A 103, 11364–11369. 10.1073/pnas.0602818103.

42. Popov, S., Popova, E., Inoue, M., Wu, Y., and Gottlinger, H. (2018). HIV-1 gag recruits PACSIN2 to promote virus spreading. Proc Natl Acad Sci U S A 115, 7093–7098. 10.1073/pnas.1801849115.

43. Hammonds, J., Chen, X., Ding, L., Fouts, T., De Vico, A., zur Megede, J., Barnett, S., and Spearman, P. (2003). Gp120 stability on HIV-1 virions and Gag-Env pseudovirions is enhanced by an uncleaved Gag core. Virology 314, 636–649. 10.1016/s0042-6822(03)00467-7.

44. Hammonds, J.E., Beeman, N., Ding, L., Takushi, S., Francis, A.C., Wang, J.J., Melikyan, G.B., and Spearman, P. (2017). Siglec-1 initiates formation of the virus-containing compartment and enhances macrophage-to-T cell transmission of HIV-1. PLoS Pathog 13, e1006181. 10.1371/journal.ppat.1006181.

45. Hammonds, J., Chen, X., Fouts, T., DeVico, A., Montefiori, D., and Spearman, P. (2005). Induction of neutralizing antibodies against human immunodeficiency virus type 1 primary isolates by Gag-Env pseudovirion immunization. J Virol 79, 14804–14814. 10.1128/JVI.79.23.14804-14814.2005.

46. Benjamini, Y., Krieger, A.M., and Yekutieli, D. (2006). Adaptive linear step-up procedures that control the false discovery rate. Biometrika 93, 491–507.

