## Supplemental Figure 1 for "Incorporation of the HIV-1 envelope glycoprotein into viral particles is regulated by the tubular recycling endosome in a cell type-specific manner"

### Supplemental Figure S1

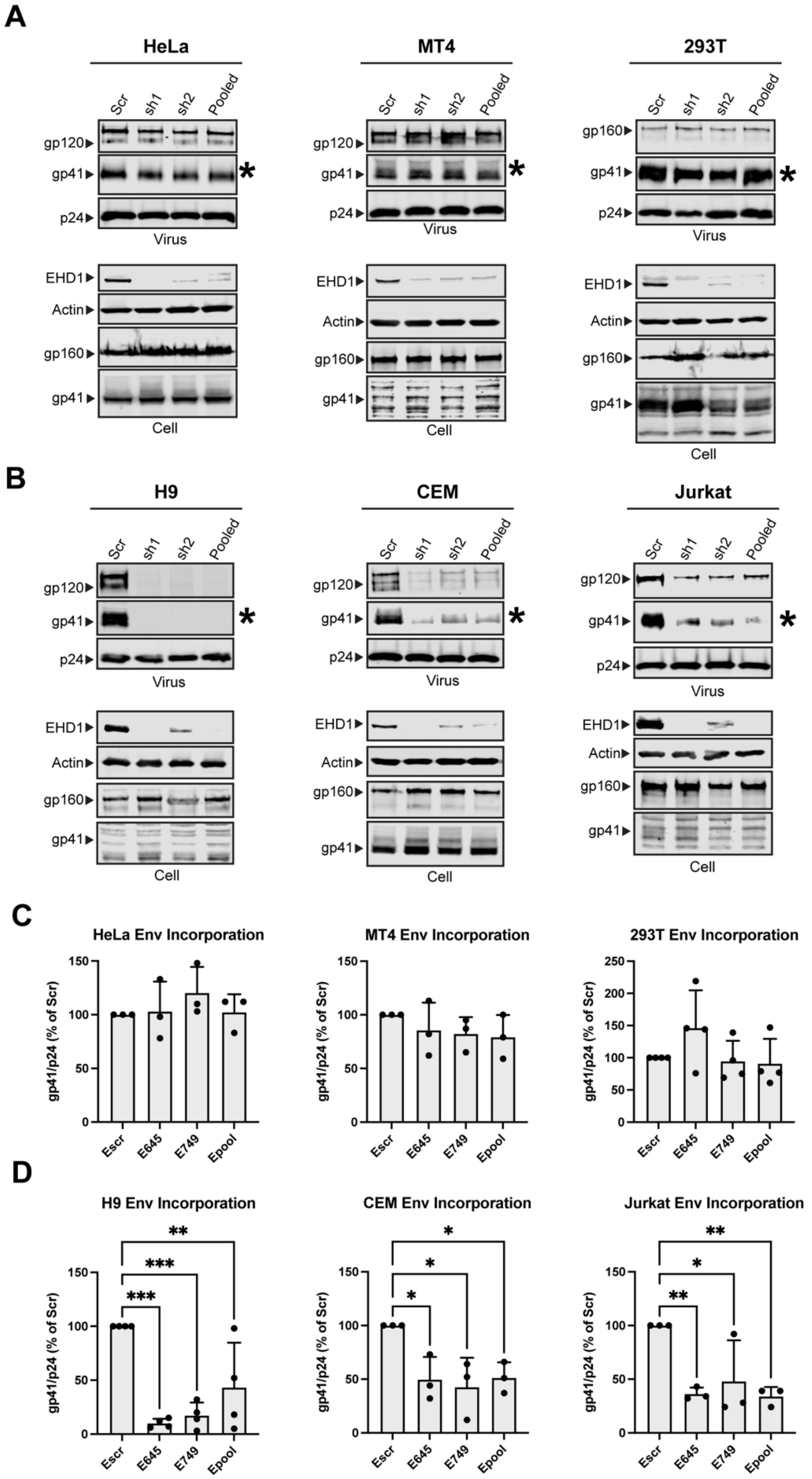

**Figure S1. Depletion of EHD1 reduces Env incorporation in a cell type-specific manner**

Viral and cellular protein content was examined following shRNA-mediated knockdown of EHD1 or treatment with scrambled (scr) shRNA. (A) Semipermissive (HeLa) cells and permissive cells MT-4, 293T displayed no noticeable defect in Env incorporation despite high efficacy of knockdown. Asterisk denotes viral gp41 lane as indicator of Env incorporation into particles. (B) Nonpermissive cell types H9, CEM, and Jurkat cells demonstrate reduced particle incorporation of Env (gp41, asterisks) following EHD1 depletion. (C) Quantitation of Env/p24 ratio, using Scr lanes as 100%, from repeated experiments; permissive/semipermissive cell panel. (D) Quantitation of Env/p24 ratio, using Scr lanes as 100%, from repeated experiments; nonpermissive cell panel.
